# Biomechanical modeling of Border Collies (*Canis familiaris*) for insights of tail and limb use during the aerial phase of jumping

**DOI:** 10.1101/2022.12.30.522334

**Authors:** Tom Rottier, Andrew K. Schulz, Katja Söhnel, Kathryn Mccarthy, Martin S. Fischer, Ardian Jusufi

## Abstract

Dogs and other members of Canidae utilize their tails for different purposes, including agile movements, such as running and jumping. In this study, we utilized motion capture biomechanical data of a border collie executing an agile rotational jump maneuver. This data created a 17-segment biomechanical model of the border collie’s (*Canis familiaris)* limb movement during agile jumps. This model was verified by comparing it to the biomechanical movement and fitting the dog’s agile task with an RMSE less than 2.5%. Using this joint model, we held specific segments constant to view their inertial impact on the dog during the aerial phase of jumping. Results suggest that the tail, hind limbs, and fore limb provides little to no inertial advantage during these rotational jump maneuvers. The tail of dogs likely does have a minimal impact on inertia, the opposite of animals like the gecko. This work could alleviate unknown biomechanical use of the tails to understand the behavioral biomechanics of lesser-known species in their ability to use their tail for rapid and taxing behaviors, including sprinting or climbing.

## Introduction

Understanding the intricacies of mammalian agile movement has long fascinated researchers across a wide range of species ranging from bipedal perturbation understanding (Andrada *et al*., 2022a, 2022b) to gait mechanics modeling of sprinting cheetahs (Patel and Braae, 2013). One powerful tool that has emerged in recent years is analytical modeling, which offers a unique avenue to delve into the mechanics and dynamics underlying the remarkable agility displayed by mammals through programs like OpenSim (Reinbolt, Seth and Delp, 2011). By constructing physical models that mimic the morphological characteristics of these animals, we can gain invaluable insights into the principles governing their agile movements and the limitations of their morphology. From the graceful leaps of cheetahs (Patel and Braae, 2013) to the precise aerial acrobatics of squirrels (Fukushima *et al*., 2021; Boulinguez-Ambroise *et al*., 2023), physical modeling provides a means to unravel the complex interplay between anatomy, biomechanics, and control strategies, shedding light on the evolution and adaptability of these remarkable creatures (Jusufi *et al*., 2010a, 2011; Shield *et al*., 2021).

An analytical 3D dynamic model of righting revealed that caudal appendage inertia suffices to reorient two lizard species from supine to prone poster during free fall via conservation of angular momentum in two squamate species(Jusufi *et al*., 2010b) whereas aerodynamic forces acting on legs predominate in invertebrates(Jusufi *et al*., 2011). Righting of cats revealed that twists and flexions of the torso are responsible for reorientation as investigated by a multisegment multijoint model (Arabyan and Tsai, 1998; Marsden and Ostrowski, 1998), although the tail is observed to rotate throughout the maneuver and was thought to be used for trim (McDonald, 1960). Based on a multibody Simulink model and the optimization analysis of the tail trajectory using a genetic algorithm, it was found that over the range of tails observed in most arboreal lizards, increasing tail length relative to body length increased the rate of torso reorientation in yaw and pitch; While lizards can use both inertial and aerodynamic forces to reorient during gliding, inertial forces appear to generate more rapid torso reorientation than utilizing aerodynamic forces (Siddall, Ibanez, *et al*., 2021). In this article, we explore the significant role of analytical modeling in unraveling the motion patterns that enable mammalian agility, showcasing its potential to enhance our understanding of both natural and robot locomotion. We propose to utilize a dynamic model constraining a quadrupedal dog to 17 segments to determine biomechanical data through added constraints.

A host of mechanical or morphological trade-offs can influence agile movement. Controlling the body’s posture is important in allowing animals to locomote through complex environments (Huang and Ahmed, 2011; Wilshin, Reeve and Spence, 2021; Biewener *et al*., 2022). For example, in enabling effective acceleration (Walter and Carrier, 2009; Hayati *et al*., 2017), agile maneuvering (Powers and Harrison, 1999; Söhnel *et al*., 2020), and turning (Powers and Harrison, 1999; Foreman, Engsberg and Foreman, 2019; Shield *et al*., 2021; Söhnel *et al*., 2021; Haagensen *et al*., 2022). In these studies, animals move in specific ways, and then analytical models are utilized to inform the changes in their body’s locomotive patterns and behavior. Experimental validation of mathematical models is enabled by robotic physical models such as during hurdle traversal in cursorial locomotion (Sehner, Fichtel and Kappeler, 2018; Siddall, Fukushima, *et al*., 2021). Moreover, regime shifts on a primate phylogeny have been reconstructed with respect to the evolution of relative tail length to identify all independent cases (Sehner, Fichtel, and Kappeler, 2018).

The tail is a unique morphological tool as in some animals the tail can have impacts of increasing the moment of inertia in mammals by upwards of 35% (Carrier, Walter and Lee, 2001), thus potentially limiting turning ability if tails are not rotated. More recently the tail as an inertial appendage has also been found in squirrels despite the tail making up a much smaller proportion of total body mass (Fukushima *et al*., 2021). However, in other animals, the tail has other uses besides biomechanical levels; animals like the snow leopard additionally utilize their long fuzzy tail for thermoregulation assistance (Mccarthy and Chapron, 2003), and other animals like the hippo utilize it for marking behavior on trees to mark off their territory (Phagan, 2019). The advantage of mechanical tail use in biomechanics is that it can be physically modeled using force and inertial equations to determine the center of mass shifts in the body, especially during agile movements like running (Shield *et al*., 2021), jumping (Fukushima *et al*., 2021), or swinging (Chang, Bertram and Lee, no date). With some species variation in tail use, in this paper, we seek to understand the limitations and capabilities of tail use during the aerial phase of an agile jumping cycle in domesticated dogs.

Canidae has been shown to exhibit various tail elevations and depressions in different movement paces, with many dogs walking with an upright tail and galloping with a tail aligned with the spinal column(Kiley-Worthington, 1976). Moreover, the tail movement reflects the limb movement (Kiley-Worthington, 1976), such passive elastic energy storage was seen on a larger scale (T-Rex) to minimize the metabolic cost of transport (van Bijlert, van Soest and Schulp, 2021). Tails have been found to assist in pitch control in leaping Velociraptors (Libby *et al*., 2012). This current study sought to design a complex biomechanical model to test the inertial capabilities of Canidae tails. The model was compared to that of a previously published study of dogs jumping (Söhnel *et al*., 2021) to examine inertial differences in tails to discuss potential behavioral implications. Therefore, our final research question is, when we view a modeled tail constrained to different motions, can we determine the biomechanical use of Canidae tails during an agile movement?

## Materials and Methods

### Dog Agile Jumping Experiments

Söhnel et al. (2021) collected the experimental data used for this study from a previous study, where full details of the experimental setup and data collection can be found. Passive markers were glued at the palpable landmarks of the Border collies for motion capturing and image analysis. A net-suit protected the long fur to cover the markers (Figure 1A). Border collies performed jumps over a hurdle while kinetics and kinematics were recorded using eight force plates (CA, Kistler Instruments AG, sampling frequency at 2 kHz) and a motion capture system (Qualisys Track Manager® software (QTM, version 2.15), 16 infrared cameras, Oqus Series 400, Qualisys, Göteborg, Sweden, sampling frequency at 400 Hz) (Figure 1B). The Border Collies performed turn jumps, as approaching the jump in a straight line, landing perpendicular to their takeoff trajectory (Figure 1B, Supplemental video). The data were processed, and the aerial phase separated before joint kinematics were calculated (Figure 1C-D).

**Figure 1.**
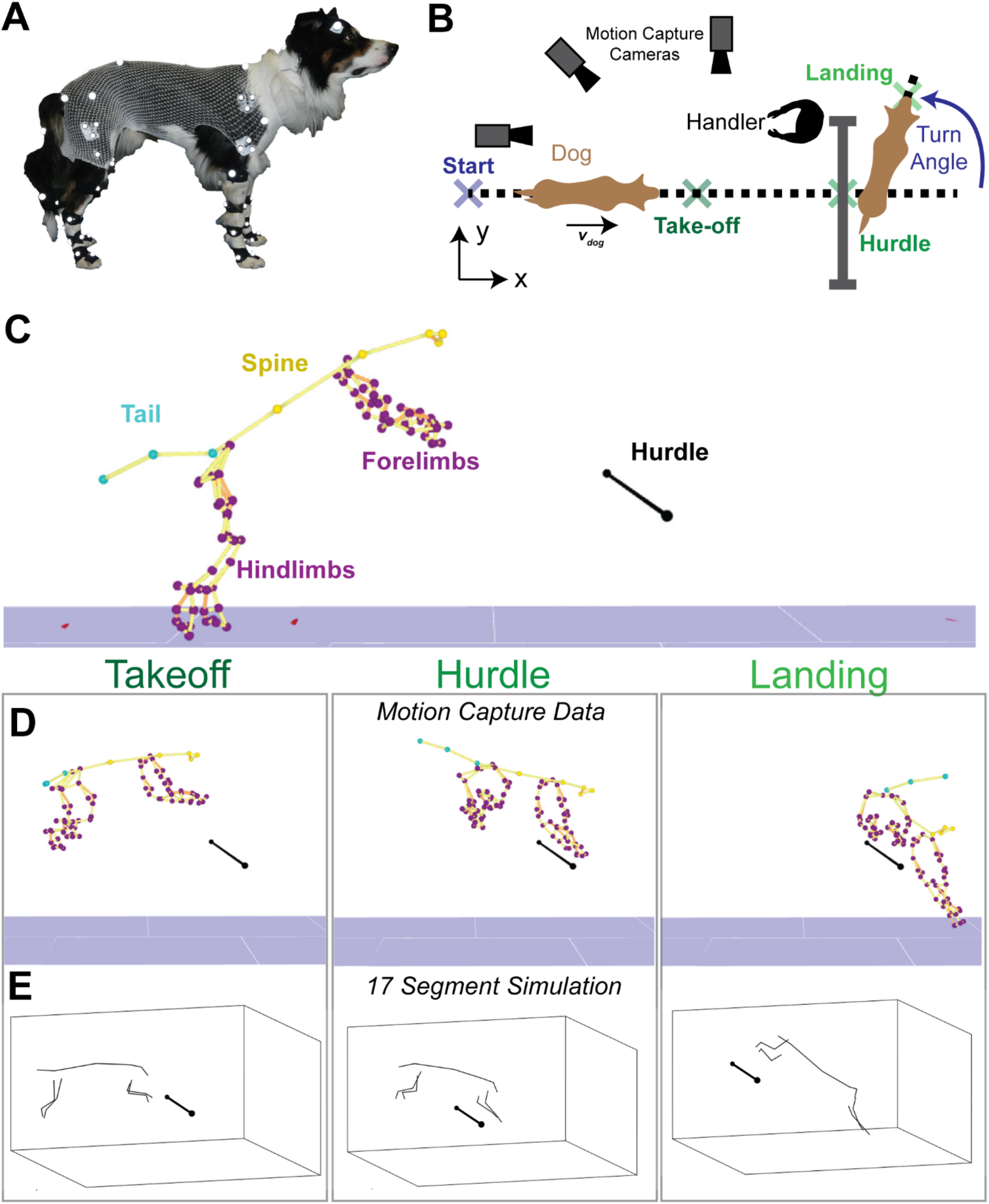
A) Experimental preparation of dog prior to motion capture with vest showing marker beads for joint and limb tracking. B) Motion capture and biomechanical arena for rotational jumping turn of dogs. C) 3D capture data of all segments from live dog prior to take-off displaying the hurdle as well as limb tracking points taken from motion capture. D) Motion tracker points extracted as a dog is jumping displaying the takeoff, hurdle, and landing phases. E) Constructed model of a jumping turn displaying the takeoff, hurdle, and landing phases from D and reconstruction of the model.

To deal with missing marker data, cubic smoothing splines were fitted to the data with any missing values removed. The splines were then evaluated over the original time to give a data set without missing data. The amount of smoothing was determined through trial and error until a good fit to the data was found through visual inspection. Forces were recorded at 2000 Hz and kinematics at 2000 Hz for the trial compared to the model. Data was not filtered for either the forces or the kinematics. For some markers, there were gaps at the beginning and end of the trial. However, they will not affect this analysis as only the middle portion of the trial, during the aerial phase, was used for further analysis. During jumping, there are three primary phases. There is the ground phase, the aerial phase, and the landing phase. The take-off was determined as follows in the transition from the ground to the aerial phase. First, the variability in the force signal was determined by calculating the average standard deviation in force across all force plates unloaded during the trial. Then the total vertical force acting on the dog was calculated by summing the vertical force across all force plates. Take-off was finally taken as the first moment the summed vertical force exceeded zero minus twice the calculated variability (as applied forces are opposing). The same approach was used to determine touchdown and thus extract the aerial phase.

The dog had several tracker dots placed along its body (Figure 1A). Overall, these tracker dots were constrained to 17 segments comprising the head, neck, upper torso, lower torso, upper limb, lower limb, and paw for each limb, and tail corresponding to the tracker beads from the study (Figure 1A). Local coordinate systems were then defined for each segment as follows: the longitudinal (z) axis was the vector joining the proximal and distal joint centers, the vertical (y) axis was the cross-product of the z-axis, and a vector joining and medial to lateral marker; the mediolateral (x) axis was then the cross-product of the y and z-axes.

For gross inertial parameters of the segments including m_limb_, I_limb_ (moment of inertia), G (COM_pos_). Initial relative values were taken from (Amit *et al*., 2009), who determined these for three breeds of dogs of differing sizes using MRI.

### Tail and Torso Morphological Reconstruction

A limitation of this study is this did not include the tail as a measurement tool. Therefore, the tail was modeled as solid cylinders with estimated radii with measured lengths from videography. The mass and torso segments of the cylinder modeled were determined from CT scans performed on a dog of the same breed (female, age 12 years, body mass 15.9kg, provided by Gifu University Japan) utilizing data from Jones et al. (2018)(Jones, Raschke and Riches, 2018). Combining these published results, we constructed the entire dog and tail scale kinematics and inertia measurements during the turning forward jump (Figure 1C-D). The orientation of the rotation taken included rotations about the z (roll), about the y (yaw), and the x (pitch) (Figure 1E-F). We proceeded to the computer and validated the 17-segment model.

### Segmented Model Creation

To model the segments during the agile forward jump of the dog in Söhnel et al., we utilized 17 segments comprising the head, neck, upper torso, and lower torso, upper, lower, and paw for each of the four limbs and tails. The modeling approach was similar to Yeadon et al., which modeled a human gymnast, so the segment approach is not similar, but the modeling framework we used in this study (Yeadon, 1990). Each upper limb joint center was displaced from the corresponding joint on the back by a constant amount that was determined by taking the mean vector between the two joint centers expressed in the upper limbs’ local coordinate system, which remained roughly constant throughout the jump. The inertial parameters of each segment were the same as those used with the experimental data.

Each segment had three degrees of freedom (DoF) which were the three Euler angles (pitch-yaw-roll rotation sequence) parameterizing the rotation of a reference frame fixed in the parent segment to one fixed in the segment. The upper torso segment defined the overall orientation of the model, with its DoF specifying the rotation of its frame from the fixed laboratory frame. Parent frames, or frames fixed in the parent segments, started from the upper torso segment and proceeded out from proximal to distal segments. The orientation angles for each segment were specified as functions of time with fitted splines to the joint angles taken from experimental data. These orientation angles, e.g., theta(t), beta(t), lambda(t), which reduced the model to an overall three DoF system which was the overall orientation of the model in the roll, pitch, and yaw directions. The functions specifying the rest of the segment’s orientation were quintic splines fitted to the experimental data, as these also gave the first and second derivatives, with respect to time, of the angles by differentiating the coefficients.

### Model Verification

The model was given initial conditions determined from the data, which were the angles and angular velocity of the upper torso at the start of the aerial phase. The kinematic and dynamic equations of motion were then integrated forward in time for the duration of the aerial phase determined from a trial to produce time histories for the three upper torso orientation angles. This trial was chosen as it was the trial that the most data could be extracted from. The total time for the aerial duration of this trial was 0.293 seconds. This model verification trial is what is reported in the comparison data.

Given the uncertainty in the model parameters, ensemble simulations were performed whereby multiple simulations were performed, each with the parameters varied slightly. For each ensemble simulation, 1000 individual simulations were performed, each with the model’s parameters randomly drawn from a uniform distribution from ± 20% of the original value. The mean and 5-95% quartiles for the simulations were then calculated.

An optimization was also performed on the parameters to minimize the mean squared (MSE) difference between the experimental data and the simulation. An evolutionary algorithm was used to vary the parameters to minimize a cost function comprising the sum of the mean squared error in the time histories for each orientation angle. Bounds were set to ± 10% of the original values. The optimization was then repeated with the bounds increased to 20% to see what effect a more considerable parameter change would have on the outcome (Figure 2B,C).

**Figure 2:**
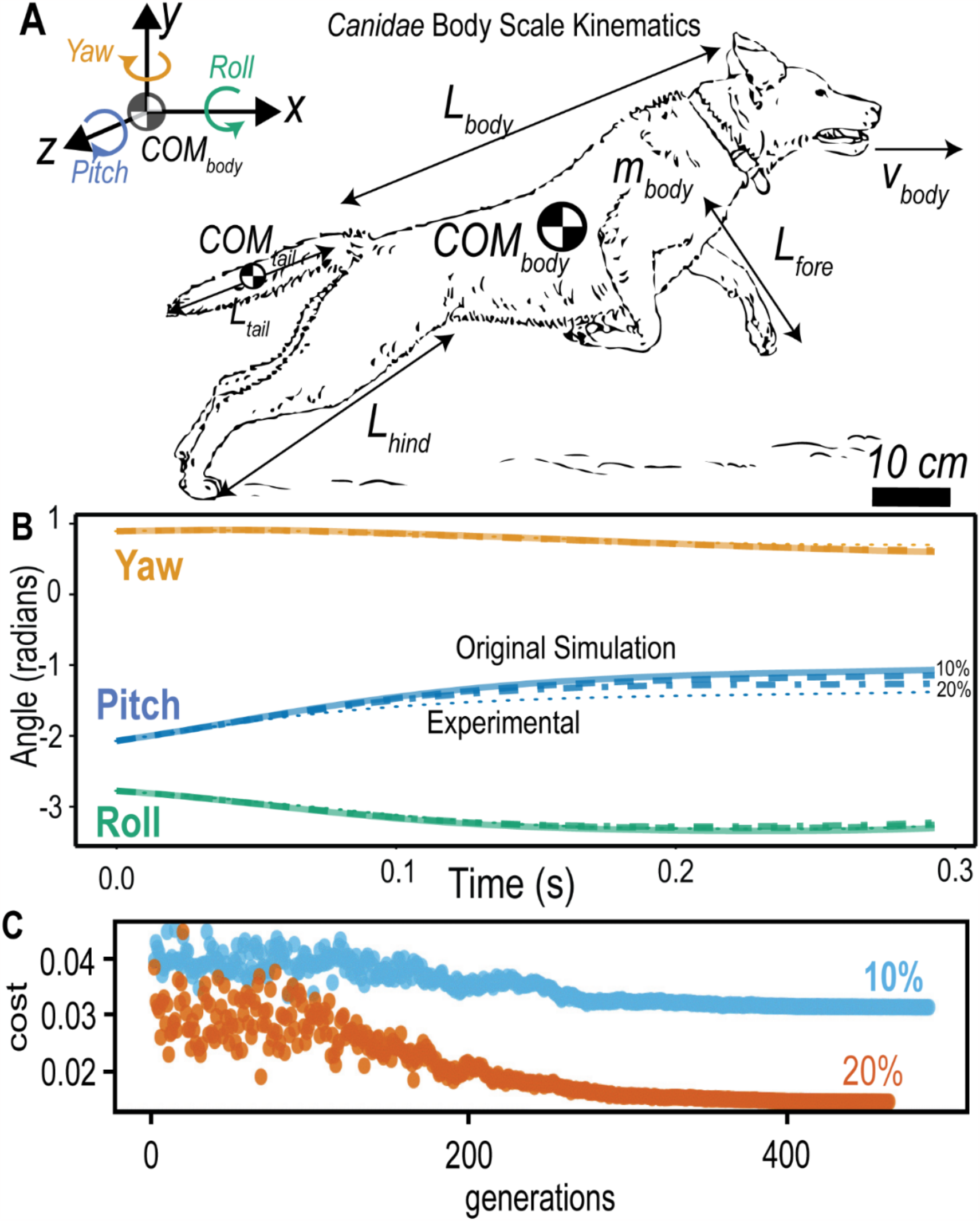
A) Body scale kinematics measurements used in the model with body rotation axis indicating Yaw in the positive y axis, roll in the positive x axis, and pitch in the positive z axis. B)Comparison of orientation angles between experimental data (dotted) and simulated (solid). Shaded region is the 5-95\% quartiles for the ensemble simulation. Illustrations from Adobe stock open source drawings. C) Cost function of generations when comparing the model and the 10 % (in blue) and 20% (in orange) to the original tracker experimental data.

**Figure 3:**
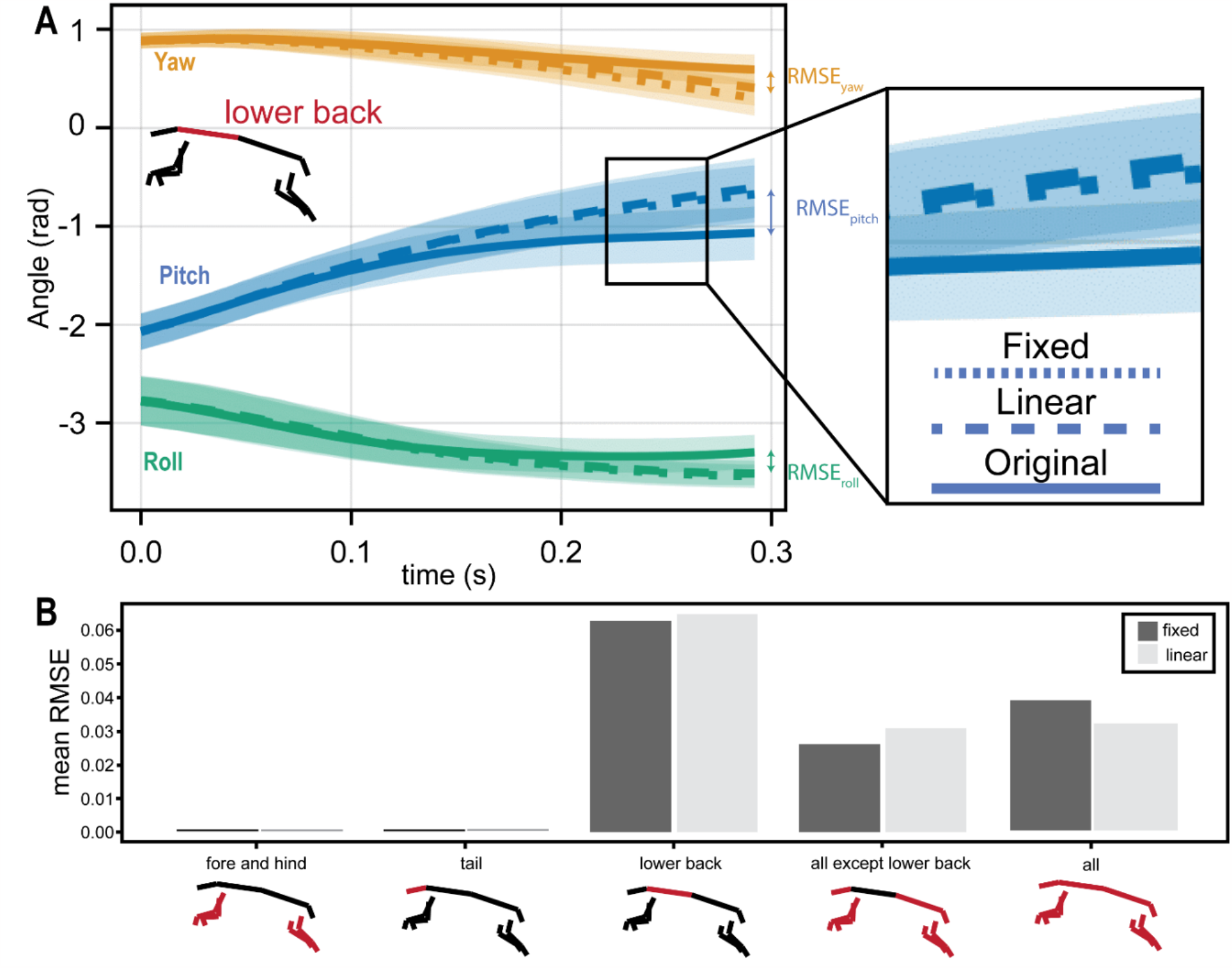
A) Angle versus time of a lower back segment displaying the change of angel versus time with the distance shown between the fixed/linear models and that of the original model. Inset displayign the difference between the three lines on the plot. B) Segments that were held constant for the tracking analysis including tail, limbs, and lower body.B) Average Root Mean Square Error (RMSE) of the different simulations with bodies held constant with showing of percentage of error between fixed and linear models. Additionally, schematics below show in red which segments of the model are held constant in the fixed and linear model.

### Constraining Joints on Model

After constructing the model and verification through uncertainty parameter verification, the ability to control the movement of each segment now meant we could explore the effects of different movements on the model’s orientation. Two alternative movement schemes were used, one where the segments were fixed at their orientation at take-off, and the orientation angles were linearly interpolated between the angles at take-off and touchdown (taken from the experimental data). We re-ran simulations with everything the same except for the altered segment orientations. Finally, we performed simulations using both schemes for all cases including: Case 1 (C1): Fore and hind limbs fixed, Case 2 (C2): Tail fixed, Case 3 (C3): Lower back fixed. Case 4 (C4): All except lower back fixed. Case 5 (C5): All segments fixed.

All simulations were performed similarly to previously published literature on biomechanical modeling given by Julia et al (Bezanson *et al*., 2015) using the SciML ecosystem(Rackauckas and Nie, 2017).

### Statistics of difference between different joint postures in border collies

We hypothesize that the dog species will have the same kinematics when specific joints are fixed in space and not given the ability to move during the agile jumping cycle. Differentiating joint mechanics of different Canidae species using the two-tailed t-test statistical method. To test the differentiation of joints being held, we used the C1-C5 case studies.

For each of these cases, pitch, roll, and yaw were calculated, as well as the original, fixed, and linear interpolated values. Statistics were calculated to determine the difference of the RMSE of all joints, lower back, and all except lower back differentiation. There was no difference between the fixed fore and hind limbs as well as the tail. To differentiate each of these cases, we did a two-tailed t-test between the fixed and linear models.

## Results

### Creating and Verification of a 17-segment Inertia Dog Model - Dog Experiments

Analysis of jumping kinematic and ground reaction forces has already been reported in Söhnel et al. (Söhnel *et al*., 2021) Therefore, we begin with creating a kinematic and inertia model of a linear dog agile jump using a 17-segment inertial model. In developing a model utilizing the parameters, inertia, and segments discussed in the methods we sought to verify the validity of this 17-segment inertial model.

The jumps analyzed were dogs landing on average a range of 75 degrees in a counter-clockwise rotation from takeoff (Figure 1B). During these jumps we analyzed the border collies jumping kinematics during the turns. We proceeded to verify the model of kinematics and inertial measurements of the dog during an agile rotational jump.

Using joint angle time histories from the experimental data of Söhnel et al. (2021) allowed us to evaluate the model by comparing the simulation outputs to the data in Figure 1F, with the shaded regions showing the 5th-95th percentile for the ensemble simulations and the solid line showing the mean. Evaluation of the model was done by minimization of the root-mean-square error between the experimental data and the 17-segment model of the border collie. Once our optimized parameters for initial angles, take-off and landing were completed we found an average RMSE for the three orientation (roll-pitch-yaw) angles was 0.0673 rad (4 degrees) (Figure 2B).

Additionally, to verify the stability of the optimized model we increased the bounds of the variables and we found that the optimization with bounds at 10% only decreased the RMSE by 0.1 degrees, or 2.5%, and the optimization with bounds at 20% decreased the RMSE by 0.8 degrees, or 20%. Most parameters changed in the same direction (increased or decreased) between the two optimizations. Given that even with optimized parameters only gave a small decrease in the cost shows that the uncertainty in the model parameters has a small effect on its outcome (Figure 2C). All optimization converged within 300 generations. With our optimized model with minimized error, we proceeded to vary different model morphological parameters of segments.

### What Joints are Inertially Relevant during Agile Dog Jumping

To test hypothesis two we constrained our model to keep specific joints and connections in the body constant during the jumping simulation. As described in the methods we tested five different case studies of segment constraints. When specifying the time histories for specific joints, or subsets of joints, the most significant effect on the orientation angles came from controlling the joint between the upper and lower torso.

This resulted in a difference in pitch, yaw, and roll angles of 0.685 rad (39.2 degrees), 0.136 rad (7.78 degrees), and 0.259 rad (14.7 degrees), respectively, at the end of the simulation between the original and with the lower body fixed. Similar but slightly smaller differences were observed when the joint angles were linearly interpolated (Figure S4A). When controlling the fore and hind limbs or tail, there was no observable difference in orientation angles (Figure S4B); in fact, when controlling all joints except the torso joint, there was only a tiny difference in the joint angles (Figure S4C).

During rotational jumping, we found that the fore-hind limbs held constant as well as the tail held constant has less than a 0.1% difference on the overall kinematics and inertia in all joint angles of the 17-segment model. It appears from our model that the dogs primarily utilize their lower back for rotational advantage.

## Discussion

Experimental data of a turn jumping border collie was used to generate a 17-segment model to test whether the tail, the limbs or the lower back have a biomechanical impact on the turn jump. A model was found that matched the experimental data well, with an RMSE und 4°. Using this model we found, unlike expected, neither the tail nor the legs have any biomechanical effect, even though *Canis* have the longest tails among *Canidae*, but the tail index, or the ratio tail to body length is the lowest.

### Biomechanics versus Behavior Across Scales

In mammals, the muscles responsible for lateral movement of the tail, flexion, and extension originate in the vertebral skeleton part of it inserted on the sacrum(Esteban *et al*., 2020). The sacrum is a crucial link in the vertebral skeleton, connecting the tail region with the presacral spine and the hind legs via the pelvis. Esteban et al. (2020) have shown that sacral morphology is related to both tail length relative to body length and body length relative to body mass.

Esteban et al. (2020) found that the shape of the sacrum in canids is related to both caudal maneuverability and spinal stabilization during locomotion. These more developed attachment areas of the muscles involved in tail maneuverability, in combination with the low tail index and the low inertial momentum, we conclude that the tail is related to social behavior (Kiley-Worthington, 1976; Hickman, 1979; Ewer, 1998; Weisbecker, Speck and Baker, 2020). Dogs seem to utilize their tails for different behavioral communication(Hecht and Horowitz, 2015), and canids have responded more positively to tail wagging as a social cue for friendliness (Reimchen and Leaver, 2008). Tails can also operate socially associated with marking behavior, like in the African Wild Dog (Parker, 2010).

At this point, we believe the border collie’s tail is primarily adapted for communication. Nevertheless, the tail moves during locomotion. As the tail is a continuum of the spine, lateral movement of the tail can be related to limb movement, where the degree of deflections tends to be related to the spinal flexibility, and flexion and extension will be in line with the curvature of the spine(Kiley-Worthington, 1976). Pretending that further study of smaller dog tail morphologies will confirm our findings, it is possible that this scaling could be nearly isometric. We also see that the tail decreases in size for larger body lengths, giving a smaller tail index.

Our results display that the tail, forelimbs, and hindlimbs do not create large differences in the inertia movement of the dogs during aerial phase of jumps. This could be due to the fact that the tail is quite small compared to the center of mass. The small amount of tail biomechanical impact on the center of mass could indicate that tail use is even less present in terrestrial locomotion than in other locomotor classifications such as arboreal, scansorial, or aquatic, where animals showed higher tail length(Esteban *et al*., 2020).

Furthermore, we were able to show that the legs also have little to no effect on the moment of inertia. The usefulness of the legs has been discussed for decades in the Falling Cat phenomenon. The upside-down falling cat can aerially righting. Loitsyansky assumed that a rapid tail movement produces torque (Polyakhov, Yushkov and Zegzhda, 2021), but this theory was rejected because (Fredrickson, 1989) showed that cats could also turn without a tail.

Further, it was also assumed that the cat rotates by changing the moment of inertia by stretching out and pulling in its legs. This model was studied by Kaufman and Essen, but this mechanism is not sufficient to explain the 180° rotation (Essén and Nordmark, 2018). However, there is hardly any obvious movement of the cat’s legs (Zhen *et al*., 2014). The most common model, on the other hand, described the modeling of the cat as a two-cylinder (the front and back halves of the cat) capable of changing their relative orientation and was advocated by many different mathematicians and physicists(Kane and Scher, 1969; Edwards, 1986; Montgomery, 1993). The multibody and multi joint model of Arabyan and Tsai 1998 suggested flexions of the spine are sufficient to explain righting.

In contrast to the tail and the limbs, we showed that the inertial impact of the lower body has a biomechanical effect on the center of mass during jumping. During turn jumping, Söhnel et al. (2021) showed that the dogs become more efficient with a well-prepared take-off phase leading in the aerial phase. The take-off should generate a high vertical, breaking, and lateral impulse by the forelimbs, lifting and shifting the torso to the new direction, followed by a higher decelerative impulse in the inner (left) hindlimb and a higher accelerative and centripetal impulse in the outer (right) hind limb, resulting in torque around the hip and a shift in the direction of the jump. Therefore, the lower body goes into the aerial phase with a turn that can be continued.

The canids sacrum morphology is related to high articulation, which is critical for an effective load transmission between the hind limbs and the presacral region of the vertebral column (Esteban *et al*., 2020) and therefore supports the animal weight during locomotion (Vleeming et al., 2012), and can be useful during the take-off phase of the jump, where the hind limbs generating the torque around the hip. The dogs can have a directional change (from straight to the left) at the beginning of the aerial phase of 25° and the end of the aerial phase of 75° turning (Söhnel et al. 2021). Esteban et al. (2020) also showed that the sacrum of canids belongs to those whose hind limbs are subjected to tremendous stress during locomotion due to a demanding locomotion behavior. As long-distance hunters, they need running endurance (Janis and Wilhelm, 1993; Figueirido et al., 2015) and high efficiency in the transmission of movement to capture prey (Janis and Wilhelm, 1993). Therefore, their skeletons must withstand stresses caused by increased muscle strain during rapid locomotion.

However, further study is needed to understand other potential biomechanical usage. We thus propose that this paper furthers the evidence that tails in the studied *Canidae* likely utilize their tails for non-agility tasks, including behavioral such as emotional indication or marking behavior.

### Limitations of Model and Agility Task

The modeling portion of this paper utilizes a complex biomechanical model of the dog and analyzes how different movements of the joints affected the outcome. Initially, the model could be evaluated against the experimentally collected data on the dogs when the same joint movements were input. The close match of the model to the data in Figure 1F, RMSE < 4◦, indicates it represented the dog quite well. Any discrepancy between the model and the data can either be due to the model itself being wrong, the parameters, or the inputs (the joint angle time histories). It is most likely the error comes from the parameters as they had to be estimated; the mode, while still making assumptions about rigidity and segment definitions, is likely complex enough, and the inputs will contain only small amounts of error. Relative values had to be taken for most segments as data for the breed of dog used is unavailable; therefore finding a dog of a similar size to tail body size values with and utilize geometric modeling was also used for some segments where relative values could not be found.

Overall, given the good match between simulation and data, these limitations do not likely affect the conclusions. We also agree that there are limitations in the agility tasks the dogs performed. This paper is primarily concluded during aerial phase of rotational jumps that dogs do not utilize their tail. The rotational jumps are a highly agile task that requires rotational movement in the body, but it is primarily a rotation in one of the axis. There are further agility tasks that could influence the movement additionally such as changing the height of the landing to be different from takeoff. Finally, the model was trained on a single dog species, and we utilized the morphological parameters of varying members of Canidae. We did not factor in morphological items like claws, or other morphological features that could be influence in the model. With these factors taken into account, we still believe that this model expansion to Canidae gives insight into tail use during agile biomechanical tasks. Expanding models in this insight can allow generating new connections between behavior-biomechanics and morphology. Additionally using a common species such as a domesticated dog allows for non-invasive studying of biomechanical movement of species that are exotic or even data-deficient on the IUCN red list.

Synthetic data models are beginning to general movement and biomechanical insight on challenging to-study species such as the Zebra. Utilizing synthetic data and mechanical models across clades allows studying of endangered and vulnerable species for greater amounts of study. There are limitations in the data access of the canid species. Many of the smaller Canidae species have only been documented through hand-drawn images, and there is almost no morphological data present at all for some of the rare and endangered species. Further study is needed on the Canidae biomechanics as well as there is no biomechanics literature present on the Canidae movement outside of varying species of domesticated dogs.

## Conclusion

In this manuscript, we describe a 17-segment kinematic model that is optimized to reduce RMSE for an agile rotational jump maneuver performed by border collie’s passive marker-based motion capturing. Through statistically differentiation, we reduce the RMSE of the model matched to the Border collie to converge at under 300 interactions. Using this minimized error model of a dog’s agile jumping we constrain subsets of the models segments to determine behavioral insights. Results suggest that the lower back is the most critical segment in inertial stability of agile jumping, while the forelimbs, hindlimbs, and tail appear to play negligible roles for conservation of inertial rotation.

## Acknowledgements

We would like to acknowledge Dr. Ryota Iwasaki, Professor at Gifu University and DVM for assistance in the morphological measurements for completing the model. We thank Dr. Robert Gillette for valuable insights. This research was supported by a Cyber Valley grant (CyVy-RF-19-08 to A.J.).

## Supplemental Figures and Information

Supplementary Video 1 Video of Dog from Experimental Trials (*Canis familiaris)*

Supplementary Video 2 Video of Dog Leaping simulation of *Canis familiaris*. https://doi.org/10.1007/s11071-014-1741-2.

